# Learning the functional landscape of microbial communities

**DOI:** 10.1101/2023.03.24.534159

**Authors:** Abigail Skwara, Karna Gowda, Mahmoud Yousef, Juan Diaz-Colunga, Arjun S. Raman, Alvaro Sanchez, Mikhail Tikhonov, Seppe Kuehn

**Affiliations:** Department of Ecology & Evolutionary Biology, Yale University, New Haven, CT 06520; Center for the Physics of Evolving Systems, The University of Chicago. Chicago, IL 60637; Department of Ecology and Evolution, The University of Chicago, Chicago, IL 60637; Department of Microbial Biotechnology, National Center for Biotechnology (CNB-CSIC), 28049 Madrid, Spain; Department of Pathology, The University of Chicago, Chicago, IL 60637; Duchossois Family Institute, The University of Chicago, Chicago, IL 60637; Department of Physics, Washington University in St. Louis, St. Louis, MO 63130

## Abstract

Microbial consortia exhibit complex functional properties in contexts ranging from soils to bioreactors to human hosts. Understanding how community composition determines emergent function is a major goal of microbial ecology. Here we address this challenge using the concept of community-function landscapes – analogs to fitness landscapes – that capture how changes in community composition alter collective function. Using datasets that represent a broad set of community functions, from production/degradation of specific compounds to biomass generation, we show that statistically-inferred landscapes quantitatively predict community functions from knowledge of strain presence or absence. Crucially, community-function landscapes allow prediction without explicit knowledge of abundance dynamics or interactions between species, and can be accurately trained using measurements from a small subset of all possible community compositions. The success of our approach arises from the fact that empirical community-function landscapes are typically not rugged, meaning that they largely lack high-order epistatic contributions that would be difficult to fit with limited data. Finally, we show this observation is generic across many ecological models, suggesting community-function landscapes can be applied broadly across many contexts. Our results open the door to the rational design of consortia without detailed knowledge of abundance dynamics or interactions.

## INTRODUCTION

Biology is the science of connecting scales of organization. From proteins to ecosystems, we are faced with the question of how variation at a lower scale of organization gives rise to changes at a higher level of organization. For example, understanding protein evolution requires learning how variation in the primary amino acid sequence determines fold and function. Similarly, at the level of the organism, genetic variation drives changes in phenotype and fitness. In both cases, interactions between constituent parts give rise to emergent functional properties.

One of the most powerful conceptual frameworks for thinking about how these functional properties emerge from components and their interactions is the notion of a landscape [1], where the height of the landscape encodes a scalar-valued function or fitness, and position on the landscape corresponds to a particular configuration of components. Landscape thinking permits us to articulate key properties of the mapping from genotypes to fitness, including the relative fitness of related genotypes [2], the extent and nature of interactions between genes [3, 4], and the dynamics of evolutionary trajectories [5–7].

Communities of microbes also exhibit emergent functional properties, from degrading complex substrates [8, 9] to resisting invasions [10–12], that arise from the constituent parts and their interactions (Fig. 1A). It is natural to ask whether the landscape concept can also be useful in these scenarios. Here we take inspiration from methods for understanding landscapes for proteins and organisms to characterize the functional landscapes of microbial communities [13].

**FIG. 1.**
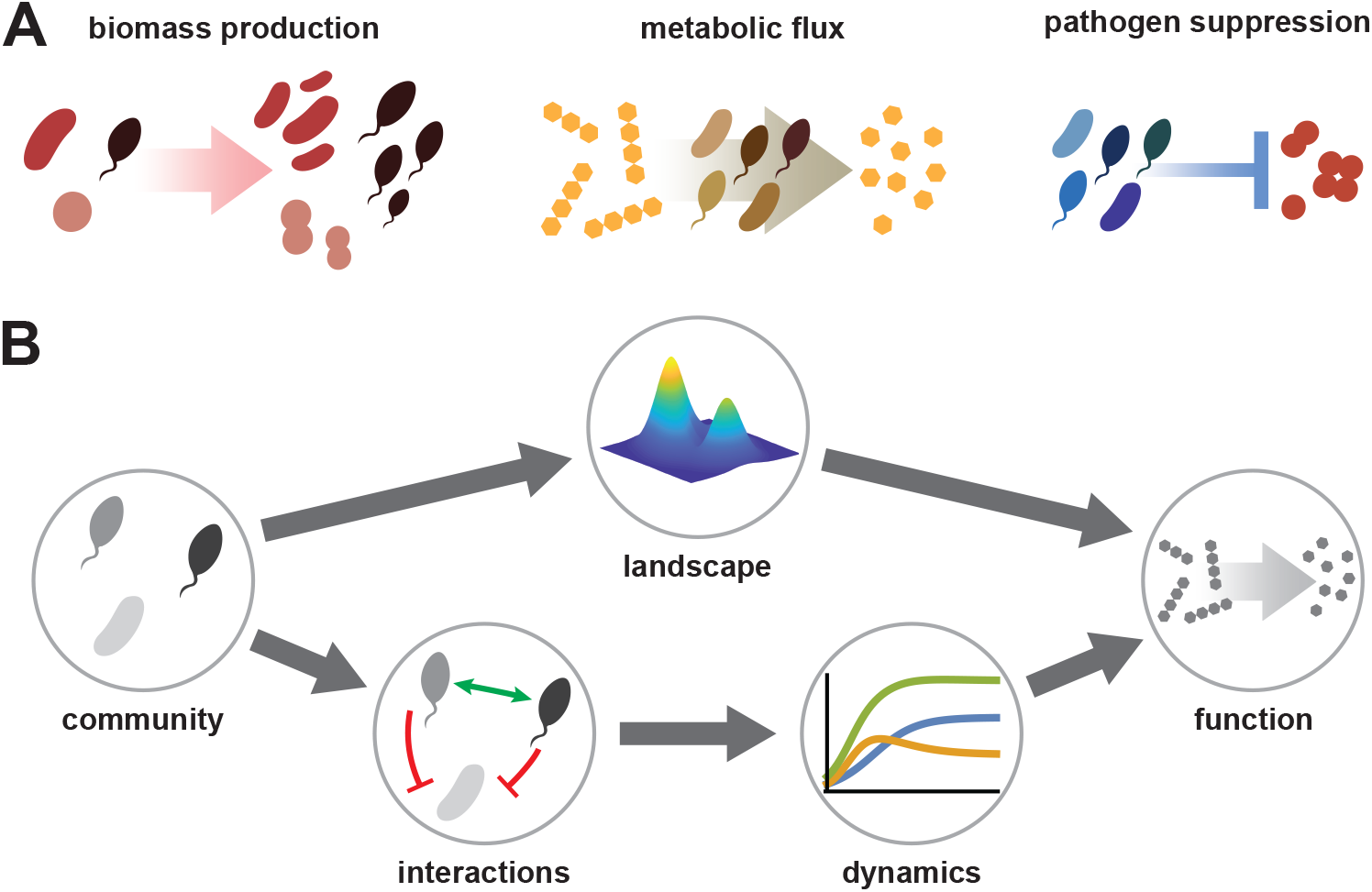
Statistically learning community-function landscapes. (**A**) Examples of microbial community functions including (left to right): production of biomass, conversion of substrate to product, and suppression of a pathogen. (**B**) Contrasting the statistical landscape view (top) of predicting community function with the dynamical view (bottom). In the dynamical view, species abundance dynamics are predicted via an ecological model, which integrates knowledge or measurements of interactions between populations. In contrast, the statistical landscape approach neglects dynamics and measures community function for a set of consortia, allowing functions for all possible community combinations to be inferred statistically.

Over the last century, it has become routine to infer landscapes in the protein and organismal context statistically [14–17]. Given that directly assaying all possible mutation combinations is typically infeasible, the statistical approach aims to approximate the landscape from a smaller number of measurements via regression [18], with the impact of interactions between mutations on fitness is quantified via nonlinear epistatic terms. It is important to note that this approach explicitly neglects the complex and often dynamical underlying processes that cause a change in fitness to arise from a mutation. For example, a statistical approach does not explicitly account for the complex physical interactions between residues that alter the function of an enzyme. Similarly, at the organismal level, this approach neglects the details of how mutations impact gene expression or life history traits. Despite this dramatic simplification, regression-based statistical approaches have been highly successful in both these contexts [2, 19, 20].

Inspired by these successes, here we take a landscape approach to quantitatively predict the emergent functions of interest in microbial communities. From this perspective, the presence and absence of species are analogous to mutations in a protein or genome, and a regression can be formulated that predicts commmunity function from species presence and absence alone. This is in contrast to most existing approaches to predicting community function, which almost exclusively seek to understand how species presence impacts abundance dynamics, and consequently function [21–24]. Here, we consider the possibility that community function can be understood without the intermediate step of predicting dynamics (Fig. 1B). We note that this approach explicitly ignores priority effects [25], multistability [26], or any other scenario when presence/absence information does not uniquely specify the community state. Nevertheless, as we will show, community-function landscapes prove remarkably predictive across a range of ecological contexts. This does not mean that the priority effects are absent; merely that, for the examples considered here, their impact on community function is, on average, weak enough that the predictive power remains high.

Implementing this approach requires measurements of community function for sets of synthetic communities constructed from libraries of taxa. Here, we utilize six existing datasets of this type, representing diverse community functional properties [21, 22, 27–29]. For all the datasets we study, we find that the functional landscape is well described by models including only additive and pairwise epistatic terms. Moreover, we find that the ruggedness of these landscapes is surprisingly low, such that the effects of species presence/absence on function are, in fact, dominated by additive terms. We support these observations computationally, showing that a regression approach succeeds in learning the community function landscape across a large class of ecological models, despite using only additive and pairwise terms, and only species presence/absence as input.

Taken together, our results show that learning the emergent properties of communities can be accomplished without a detailed understanding of the interactions between taxa or their abundance dynamics. Our findings enable a powerful conceptual framework for predicting community functional properties, from invasion resistance to biotechnological applications.

## RESULTS

### Learning community-function landscapes via regression

We first formulated a statistical approach to fitting community-function landscapes using datasets that comprise measured values of microbial community functions for a set of defined species combinations. In each of these experiments, a pool of defined species was used to combinatorially construct communities. Each community was then incubated, typically for a defined period of time, and then a functional property of interest was assayed. For a pool of *N* total species, there are 2^*N*^ *−* 1 possible species combinations, and measuring all possible combinations is frequently intractable. The first goal of our investigation was to ask whether we can predict the function of all 2^*N*^ *−* 1 communities by fitting a statistical model to a small subset (≪ 2^*N*^ *−* 1) of possible measurements. The resulting model would provide a global picture of the community-function landscape.

We formulated this problem as a linear regression of the following form:

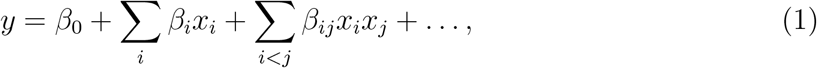

where *y* is the scalar-valued function, and *x*_*i*_ represents the presence or absence of species *i* in that community. The coefficient *β*_*i*_ is the additive effect of adding species *i* to the community, and *β*_*ij*_ is analogous to the effect of pairwise epistasis in genetic fitness landscapes, which measures the impact beyond individual additive effects of adding both species *i* and *j*. The ellipses denote higher-order epistasis terms, e.g., three-way epistatic terms captured by third-order polynomials, and so on.

We took the convention that *x*_*i*_ = 1 if species *i* is present in a community, and *x*_*i*_ = *−*1 if that species is absent. We denoted absence using *−*1 instead of 0 in order to simplify interpretation of the regression coefficients (see discussion in Ref. [30]). In brief, using *x*_*i*_ = *±*1 allows us to interpret *β*_*i*_ as the average effect of adding strain *i* to community function, where the average is taken over multiple community compositions. Similarly, the pairwise coefficient *β*_*ij*_ captures the average epistatic effect of species *i* and *j* together across many consortia. Moreover, this convention has some convenient mathematical properties (corresponding to the Fourier expansion of the landscape) that make it easier to quantify how much variation in the measured function is captured by additive, pairwise, and higher-order terms [30, 31].

We considered regressions truncated at first, second, and third orders. Many of the datasets we utilized in our investigation sampled a number of community configurations that is comparable to the total number of coefficients to be inferred. To mitigate the risk of overfitting, we employed *L*_1_-regularized regression (LASSO) [32], using a cross-validation procedure to estimate the regularization hyperparameter (Methods). To assess out-of-sample generalization error, we applied an additional leave-one-out cross-validation scheme, in which each data point was iteratively left out-of-sample, and the model was fit to all remaining data points, allowing an out-of-sample prediction for each distinct experimental community (Methods).

### Community function is predictable from strain presence/absence

We compiled six datasets in which synthetic bacterial communities were assembled from a pool of species. These datasets represent a broad spectrum of community functions: Clark *et al*. [22] measured the production of the short-chain fatty acid butyrate (Fig. 2A); Langenheder *et al*. [27] and Sanchez-Gorostiaga *et al*. [21] measured the breakdown of mono- and polysaccharides, respectively (Fig. 2B,C); Diaz-Colunga *et al*. [28] measured the total production of iron-scavenging siderophores (Fig. 2D). In addition, we considered biomass-related community functions: Diaz-Colunga *et al*. [28] measured total community biomass (Fig. 2E), and Kehe *et al*. [29] measured the abundance of a single target species (Fig. 2F). Details about the size and species pools for each dataset are given in Table S1.

**FIG. 2.**
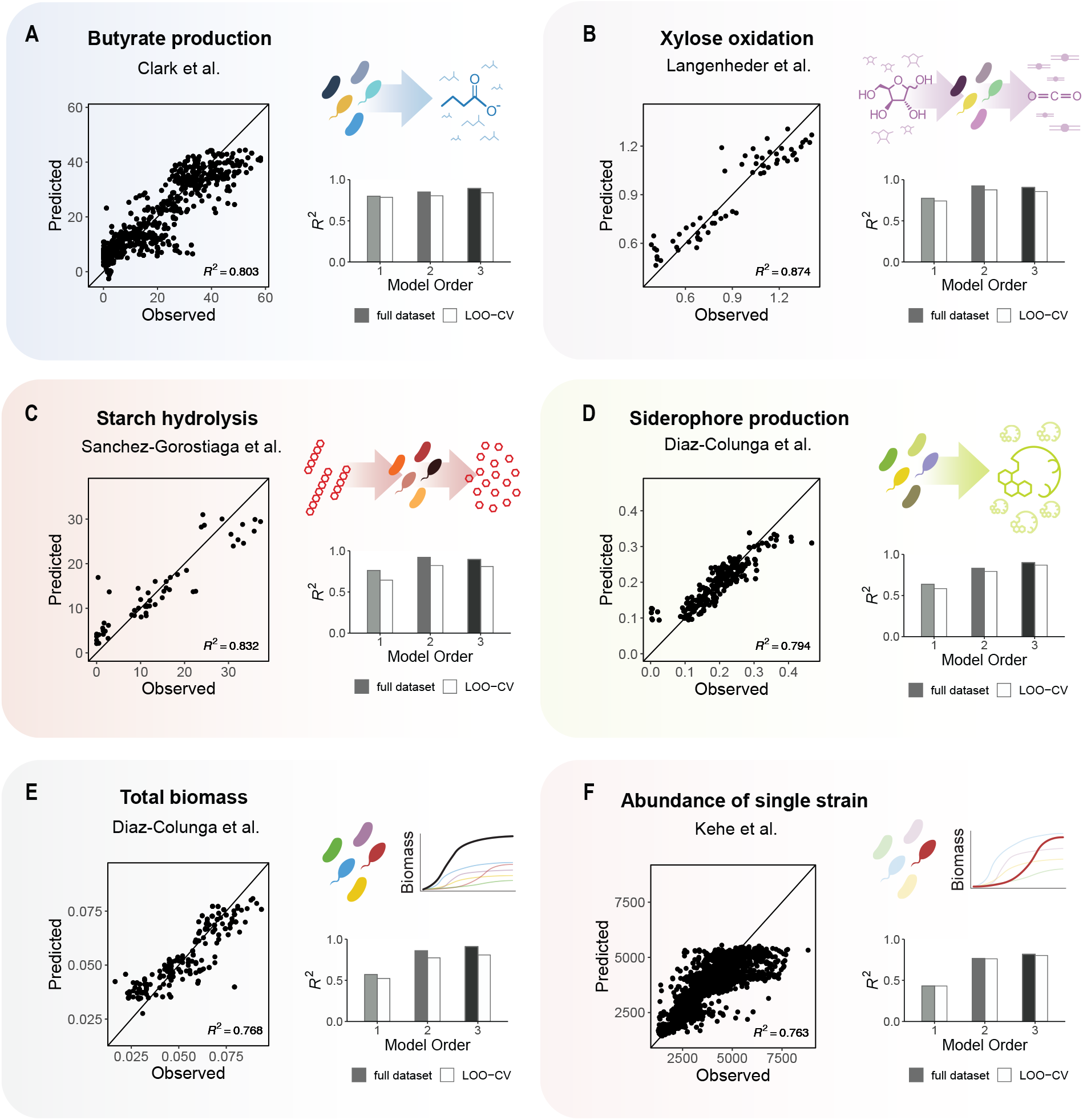
Community function is predictable from strain presence/absence in empirical datasets. For each dataset, regularized linear regressions were performed using models (Eqn. 1) truncated at the first, second, and third order. Results of model fits either using all experimental data or using a systematic leave-one-out cross validation approach are shown, with the quality of fit (*R*^2^) reported in bar plots for each model. Scatter plots show out-of-sample prediction values for the second-order regression fits obtained via the leave-one-out cross validation procedure. Analyses are shown for datasets by (**A**) Clark *et al*. [22], (**B**) Langenheder *et al*. [27], (**C**) Sanchez-Gorostiaga *et al*. [21], (**D**–**E**) Diaz-Colunga *et al*. [28], and (**F**) Kehe *et al*. [29].

In each dataset, community function can be defined as a measurable scalar quantity, e.g., the concentration of a compound or a measurement of biomass at a given point in time. Therefore we fit models of the form shown in Eqn. 1. We investigated truncating the model at successively higher orders of epistatic terms in order to determine what degree of model complexity is needed to accurately predict community function. The quality of the fits are shown in Fig. 2 (bar plots) for increasingly complex models, with shaded bars showing in-sample *R*^2^ and white bars showing out-of-sample *R*^2^ (Methods); scatter plots visualize the quality of second-order out-of-sample model predictions.

Remarkably, across all six datasets studied here, models of first or second order provide high-quality predictions (*R*^2^ *∼* 0.8). In most cases, additive models alone (e.g., *y* = *β*_0_ + Σ _*i*_ *β*_*i*_*x*_*i*_) already have strong predictive power (*R*^2^ *>* 0.5), and adding second-order terms yielded only marginal improvement. The exception is the dataset by Kehe *et al*. quantifying the abundance of an individual strain in the community (Fig. 2F), where second-order terms significantly improved predictions. We note that residuals of observed versus predicted values demonstrate similar patterns of heteroskedasticity across datasets, where high-function communities tend to be underestimated, and low-function communities tend to be be overestimated (scatter plots, Fig. 2). It is likely that these patterns are a consequence of bias induced by regularization [33].

These results indicate that, across a range of contexts, the emergent functional properties of microbial communities can be predicted using simply knowledge of which strains are present. We note that for the dataset by Clark *et al*. (Fig. 2A), the authors predicted butyrate production using a complex model that parameterized interactions and abundance dynamics. Our predictions using strain presence/absence are similar to those obtained using a complex dynamical model [22], suggesting that detailed dissections of community dynamics are not always necessary to make reasonably good predictions of community function.

### Empirical community-function landscapes are not rugged

We demonstrated that statistical models of species presence/absence can predict microbial community functions to a surprising degree of accuracy. In particular, simple models, containing only additive and/or pairwise epistatic terms, explained the vast majority of the variation in the data (Fig. 2). Our statistical approach represents a strategy for approximating the empirical community-function landscapes for these datasets. We wanted to gain intuition for why these regressions appeared to be so successful. To do this we sought to quantify the ruggedness of community-function landscapes.

In an evolutionary context, the ruggedness of a fitness landscape dictates the number of local fitness optima. Landscapes that are rugged, containing many large peaks, have important implications for the predictability of evolution [5]. In a community context, very rugged landscapes are expected to be much harder to approximate globally using regression methods such as those used here, simply because ruggedness arises from a substantial number of strong high-order epistatic terms. High-order terms are challenging to learn statistically due to the explosive combinatorial increase in the number of terms as model order increases. Thus, we expect that rugged landscapes will be difficult to learn statistically, while non-rugged ones will be straightforward to approximate using low-order models.

We quantified ruggedness using two complementary approaches. First, we made use of the combinatorially complete dataset from Langenheder *et al*. [27], in which xylose oxidation was measured for all 2^6^ *−* 1 = 63 species combinations that can be formed from a 6-species pool.

This combinatorially complete dataset allowed us to compute the coefficients of the *exact* full-order empirical landscape, which is a model of the form Eqn. 1 that includes epistatic terms of all possible orders (i.e., up to sixth order). Using this exactly-inferred landscape, we generated a Fourier amplitude spectrum (Methods), which is a decomposition that reflects the total variance of the landscape that is captured by terms in each order [30, 31]. This spectrum, shown in Fig. 3A (red line), indicates that ∼78% and ∼16% of the variance in the landscape is explained by first and second-order terms, respectively, leaving ∼6% of variance remaining for higher-order terms. In other words, this exact community-function landscape displays a low degree of ruggedness, as it is dominated by low-order terms, and is largely free of consequential higher-order terms.

**FIG. 3.**
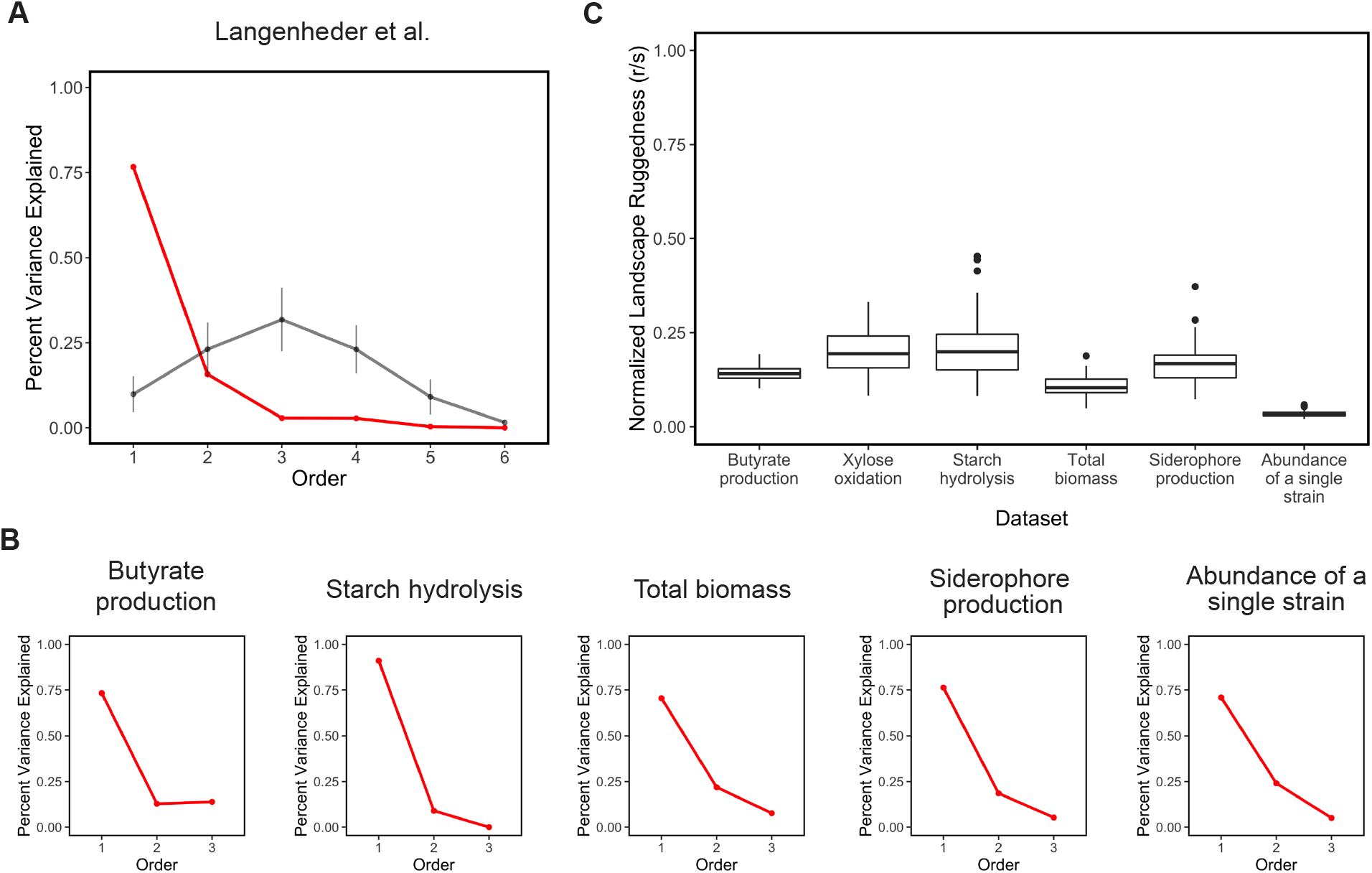
Empirical community-function landscapes are not rugged. (**A**) Normalized Fourier amplitude spectrum obtained from the combinatorially complete landscape from Langenheder et al [27]. Normalized amplitude values correspond to the fraction of total landscape variance that is captured by terms at each order. The empirical amplitude spectrum (red) demonstrates that landscape variance is primarily explained by first-order terms, with higher-order terms explaining decreasing fractions of the variance. This denotes a low degree of ruggedness in the empirical landscape. Comparisons with spectra obtained from 100 randomized landscapes (black) show that first-order terms explain a relatively small fraction of total variance, reflecting the ruggedness of the randomized landscapes. (**B**) For each additional dataset that is not combinatorially complete, the fitted coefficients of the regularized third-order linear regression are used to infer the normalized amplitude spectrum at first through third order. As in panel **A**, the amplitudes of first order terms are notably larger than amplitudes of second and third orders, indicating a lack of ruggedness. (**C**) For each dataset, normalized *r/s* values are computed by calculating the ratio of empirical *r/s* values on original landscapes to the *r/s* value of 100 randomized landscapes. The normalized *r/s* values for all datasets are notably smaller than 1, again indicating a low degree of ruggedness in empirical landscapes compared to their randomized counterparts.

To assess the significance of this result, we performed a randomization test by computationally shuffling the assignments between function measurements and community compositions. For 100 such randomizations, we inferred exact landscape coefficients and computed the Fourier amplitude spectra. The results are plotted in Fig. 3A (black line). In the randomized landscapes, we found that terms of third-order were most important, and additive terms alone captured only 10% of the variance; the peak at third order arises from the fact that there are combinatorially more terms possible at this order than any other. We concluded that a lack of ruggedness is not spurious but rather a distinctly non-random structural feature of this landscape.

We performed similar analyses for the remaining five datasets to estimate the relative importance of coefficients of different orders. Although these datasets are not combinatorially-complete, and therefore do not permit the inference of the exact amplitude spectra, we inferred a truncated amplitude spectra via third-order regression. We found that in each case, the dominant amplitude corresponds to additive coefficients, with pairwise coefficients typically the next most-important (Fig. 3B). The large fraction of variance explained by additive and pairwise coefficients across these cases again indicates a low degree of ruggedness.

Second, for each dataset we investigated the roughness/slope ratio (*r/s*) [34, 35], a metric for ruggedness that quantifies how well a landscape is fit by a purely additive model. Explicitly, roughness *r* is computed by fitting a model with additive terms only (e.g., *y* = *β*_0_ + Σ _*i*_ *β*_*i*_*x*_*i*_) and computing a residual to this fit. The ruggedness (*r*) is then the standard deviation of these residuals. Slope *s* is defined as the mean (absolute) value of the additive coefficients *β*_*i*_. The ratio *r/s* then represents the typical size of additive model error relative to the typical size of an additive term. Large values of this ratio mean that the landscape is not well approximated by an additive model, indicating a high degree of ruggedness.

To make a well-defined comparison between all datasets, we computed normalized *r/s* values, defined as the ratio between *r/s* on the original dataset and *r/s* for 100 randomized landscapes (Methods). Randomized landscapes served here as a natural high-ruggedness comparison. We found that this ratio was consistently ≪ 1, indicating that true landscapes are much less rugged than comparable random landscapes (Fig. 3C).

We concluded that, across a diverse range of community functions, empirical landscapes possess a low degree of ruggedness. Crucially, this lack of ruggedness underlies the success of our statistical approach to predicting community function from species presence/absence. In short, a low degree of ruggedness corresponds to the dominance of low-order (i.e., additive and pairwise epistatic) terms, and a notable absence of higher-order epistatic terms. This enables low-order statistical models to faithfully parameterize empirical landscapes and thereby accurately predict community functions.

### Ecological models indicate both optimism and caution for inferring community-function landscapes

Finally, having demonstrated that low-order statistical models can predict community functions in empirical datasets due to the non-rugged nature of the underlying landscapes, we turned to ecological models to further probe the conditions under which these results are likely to hold. We also attempted to identify scenarios where community-function landscapes exhibit a degree of ruggedness that would make them challenging to infer statistically with low-order models.

We first sought to interrogate synthetically-generated community-function landscapes using the generalized Lotka-Volterra (gLV) model, with total abundance (biomass) serving as the function of interest. The gLV model has many variants that differ in assumptions regarding the structure of randomly-generated species interactions, but as recently argued by Barbier *et al*. [36], a large class of such variants can be reduced to a four-parameter “reference model.” We adopted this reference model for our analyses (Methods), focusing on a sweep of the two key parameters describing interaction strength (*µ*) and interaction variability (*σ*).

For each combination of *µ* and *σ*, we performed 10 trials of generating a random pool of *N* = 10 species (a pool small enough to evaluate the combinatorially complete set of all possible communities), and computed the exact community-function landscape (mimicking our procedure for the Langenheder *et al*. data in Fig. 3A). The fraction of variance explained by the first two orders of terms, equivalent to the predictive power (*R*^2^) of the second-order approximation of the landscape, is shown in Fig. 4A. These values are averaged over the 10 trials at each point in the *µ*-*σ* plane. Note that as interaction variability *σ* becomes large, the Lotka-Volterra model becomes unstable, causing species abundances to diverge [37]. In our simulations, any points in *µ*-*σ* plane that encountered divergences in more than 5 trials are indicated in grey (Fig. 4A). Additional details about these simulations are described in Methods.

**FIG. 4.**
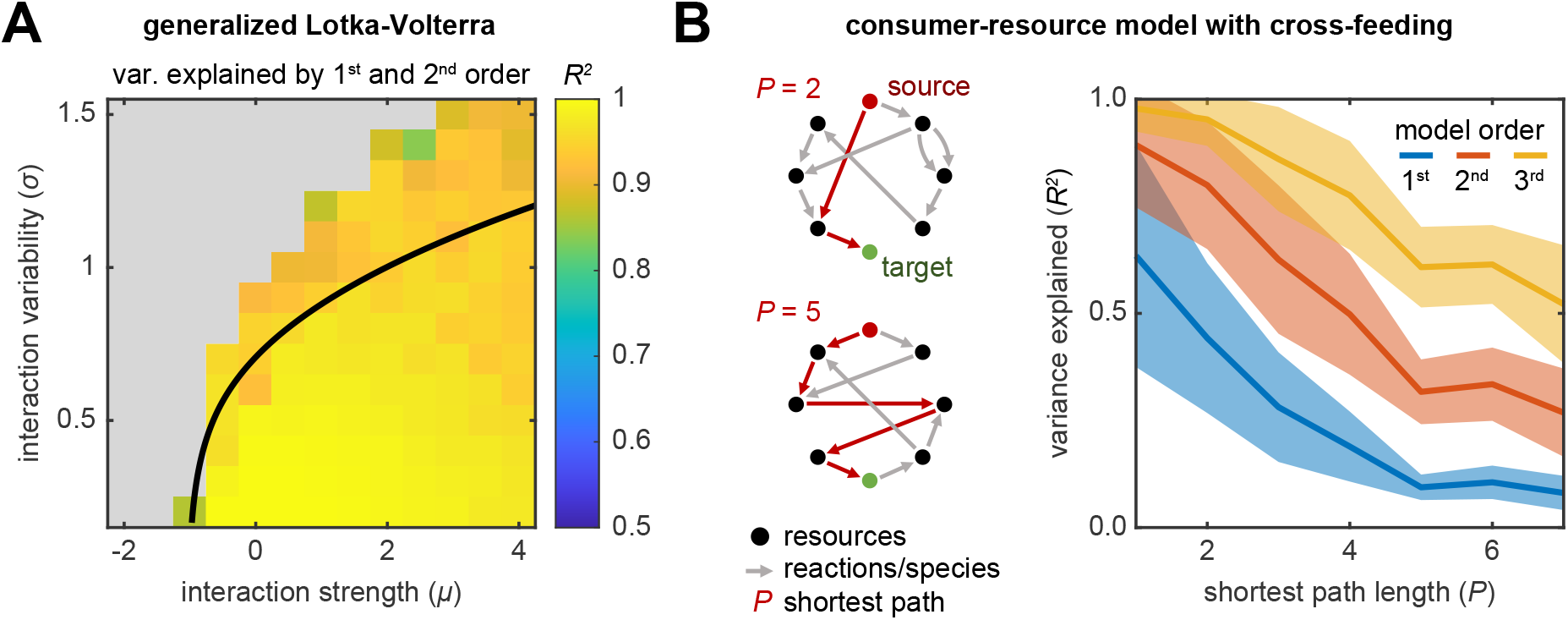
Ecological models indicate both optimism and caution for inferring community-function landscapes. (**A**) The generalized Lotka-Volterra (gLV) model was used to generate combinatorially-complete synthetic community-function landscapes, taking total abundance as the function of interest. These landscapes were then fit to Eqn. 1 exactly, with the variance explained by combined first and second order terms shown (*R*^2^). Landscapes were generated using an *N* = 10 species pool, and two parameters controlling the structure of randomly-drawn interactions were varied: an interaction-strength parameter *µ*, and an interaction variability parameter *σ*. 10 landscape trials were generated over a grid of points in the *µ*-*σ* plane, and *R*^2^ values averaged over these trials are shown. Note that as interaction variability *σ* becomes large, the model becomes unstable, causing species abundances to diverge. In our simulations, any points in *µ*-*σ* plane that encountered divergences in more than 5 landscape trials are indicated in grey. The black curve shows an analytical prediction (computed in the *N → ∞* limit [37]) for the stability boundary of the gLV model, beyond which species abundances will typically diverge. (**B**) A consumer-resource model (CRM) was used to generate synthetic community-function landscapes with randomly-generated cross-feeding networks (schematic). One resource was supplied externally (“source”), while the function of the community was defined to be the concentration of a different resource (“target”). These combinatorially-complete landscapes were fit exactly, with the variance explained by models truncated at first, second, and third-order shown (*R*^2^). Cross-feeding networks, comprising *N* = 10 species and *L* = 8 resources, were generated in order to vary the length of the shortest path (*P*) connecting the source resource to the target. The variance explained by first through third-order models are shown as a function of *P*.

We found that across the entire range of parameters for which the dynamics of the model are stable, second-order regression provides excellent fits (*R*^2^ ≳ 0.9) to gLV community-function landscapes (Fig. 4A; the non-gray region is all yellow). In the reference model, positive *µ* corresponds to interactions that are competitive on average, while larger values of *σ* correspond to a greater degree of ecological heterogeneity. High quality fits across a broad sweep of the *µ*-*σ* parameter space provides strong evidence that our findings from six empirical datasets applies to a wide range of ecological scenarios.

Despite the evident simplicity of community-function landscapes in both empirical datasets and in the gLV model, it is possible to conceive of scenarios for which a low-order approximation will be a poor one. A simple case in point is the example of a nutrient *X*_1_ that is broken down through a chain of *L* reactions following a linear pathway 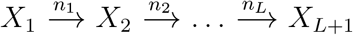, where each reaction is performed by a specialized community member *n*_*i*_. If *X*_1_ is the only nutrient supplied to the consortium, and the function of interest is the concentration of the end product *X*_*L*+1_, then the concentration of *X*_*L*+1_ will be non-zero if and only if all species *n*_1_, … *n*_*L*_ are present. Mathematically, if presence or absence were denoted as *x*_*i*_ = 0 or 1, respectively, such a landscape would be described by a single *L*th-order term: *y* = *β*_1…*L*_ *x*_1_*x*_2_ … *x*_*L*_. Under the convention used here, where presence or absence is denoted by *x*_*i*_ = 1 or *−*1, respectively, this corresponds to interactions of all orders being equally strong. In either case, it is clear that no low-order approximation could adequately capture this landscape.

To build on the intuition from this thought experiment, we used a consumer-resource model (CRM) to generate synthetic landscapes with *N* = 10 species competing for *L* = 8 nutrients. Only one resource was supplied externally (“source”, Fig. 4B, red dot), while the function of the community was defined to be the concentration of one of the other resources (“target”, Fig. 4B, green dot). In each trial, we constructed a random cross-feeding network for a pool of *N* = 10 species, with each species capable of converting one randomly chosen resource into another. We then computed the complete functional landscape, as above for the gLV, by simulating all possible subsets of this 10-species pool. Based on the intuition above, we expected the predictive power of low-order approximations to correlate with the length of the shortest path (*P*) connecting the source resource to the target (see schematic, Fig. 4B). As in the thought experiment, cross-feeding networks with long paths from source to target (i.e., large *P*) are expected to have rugged landscapes that would be poorly characterized by a low-order model. Simulation details are described in the Methods.

The predictive power (*R*^2^) of low-order models fit to these synthetic CRM landscapes is shown in Fig. 4B as a function of the shortest chain length *P*. We observed that, while successful in cross-feeding networks with small *P*, low-order models were increasingly challenged as *P* increases, with *R*^2^ dropping below 0.5 for second-order models beyond *P* = 4. Increasing model order from additive-only to third-order substantially improved predictive power, particularly at large *P*, consistent with the idea that increasing *P* increases the prevalence of higher-order terms. These results illustrate that in ecological scenarios with long chains of trophic dependencies, or other situations when function is strongly contingent on the presence of multiple species with little functional redundancy [38], it is possible that low-order approximations of community-function landscapes will fail to make accurate predictions.

## DISCUSSION

The key result of our study is the demonstration that emergent functional properties of microbial communities can be predicted simply from knowledge of which strains are present or absent. Remarkably, in analogy to fitness landscapes describing proteins and organisms, we showed that regressions can quantitatively describe empirical landscapes for a wide range of community functions. We found that the success of these regressions derives from the fact that the underlying landscapes are not rugged, allowing the majority of variation in function to be captured by additive and pairwise terms in the regression.

Our simulations of generalized Lotka-Volterra and consumer-resource models demonstrated that we can expect low-order approximations of landscapes to work well across a range of ecological contexts. We identified a clear exception, however, in functions that are strongly contingent on the presence of multiple highly specialized community members with little functional redundancy. One might therefore expect that the low-order landscape approximations might encounter challenges, for example, in systems with long linear chains of reactions, such as those present in anaerobic digesters [39] and Winogradsky columns [40]. An important direction to be addressed in future work is a systematic analysis of ecological mechanisms that either enable or impede the performance of approximations of community-function landscapes. A better understanding of these mechanisms would provide a more principled view of the community functional properties amenable to our statistical approach. Our results complement previous studies demonstrating that community-function landscapes follow patterns of global epistasis [28]. These patterns were first discovered in the context of organismal fitness landscapes [41–43]. Global epistasis refers not to the impacts of individual epistatic terms, but rather properties (e.g., diminishing returns) that emerge from the collective impact of many epistatic contributions. In the global epistasis approach to predicting community function, the effect of adding a strain follows a simple linear relationship that depends only on the identity of the strain and the value of a the function in the “background” community. It remains an important exercise to connect the concept of global epistasis to the regression approach explored here, as the two approaches may provide complementary value in different ecological scenarios.

Perhaps the greatest downstream impact of our study is the possibility of using statistically-inferred landscapes to rationally design communities with desired functional properties. Because community-function landscapes can be approximated well by sampling only a subset of all possible species combinations, it is conceivable that even large synthetic consortia with predefined functions can be designed and optimized computationally given only a small number of measurements. Across the datasets considered here, our models were consistently able to identify the communities with the highest functional output, even when these communities were left out of sample. These high-function communities are accurately identified despite the heteroskedasticity of the model fits (Fig. 2), which is likely due to a bias induced by regularization [44]. Given the importance of regularization for training predictive low-order models of landscapes using limited data, an impactful direction for future study will be the investigation of regularization approaches (e.g., weighting schemes) that minimize prediction error for high-function communities.

Finally, our approach did not require specialized knowledge or measurement of microscopic system properties, and is thus readily portable across contexts and functions. We believe this represents a major advance relative to alternative design approaches, which utilized detailed models of metabolism [45] or ecological interactions [22]. The approach we have demonstrated could enable the design of communities in a wide range of contexts, from biotechnology to therapeutics.

## ACKNOWLEDGMENTS

SK, KG, and MY acknowledge funding from the National Science Foundation (EF-2025293, MCB-2117477). AS acknowledges support from the Spanish Ministry of Science and Innovation under project PID2021-125478NA-100. SK acknowledges useful discussions with the members of the Center for the Physics of Evolving Systems at the University of Chicago.

## DATA AVAILABILITY

Data analyzed here is either available from the original studies or in the following repository: https://github.com/abbyskwara2/regression_on_landscapes.

## METHODS

### Collection and preprocessing of datasets

Datasets were compiled from six experimental efforts to measure various community functions in defined synthetic microbial consortia. Details about these datasets and references listed in Table S1.

Total community biomass data collected by Diaz-Colunga *et al*. [28] were not published in the original manuscript, though detailed methods describing these experiments and measurements are described therein. Briefly, biomass was estimated in defined communities by measuring OD600 (Multiskan FC) of a 200 µL homogenized cell suspension after 48 h of growth in M9 media with 0.2% w/v citrate as the only carbon source.

Data from Kehe *et al*. [29] were generated using a microwell array approach, in which each community was assembled by randomly grouping nanoliter droplets of defined species composition into 2–19 droplet combinations. Due to this stochastic assembly, initial species abundances varied beyond binary presence/absence: for example, a three-droplet combination containing two droplets of species A and one droplet of species B will have different initial abundances than the combination of one droplet of species A and two droplets of species B. Since the formulation of community-function landscapes in Eqn. 1 operates on binary species presence/absence, this variation in initial abundances was ignored, and a strain was considered to be present in a community if it had positive initial abundance.

In cases where datasets contained experimental replicates, the mean over replicates was taken in order to obtain a single community function value for each unique species combination.

### Statistical inference of landscapes via regression

Community-function landscapes were approximated for empirical datasets by fitting equations of the form Eqn. 1 truncated at first, second, and third order via LASSO regression [32]. 10-fold cross-validation was used to estimate the regularization hyperparameter. All models were fit using the package glmnet in R version 4.1.2.

Two different training strategies were employed. First, all available data points were used to fit landscape models. The coefficients of determination (*R*^2^) for these fits are shown in grey bars in Fig. 2. Second, to obtain an estimate of out-of-sample model accuracy, a leave-one-out (LOO) procedure was employed. Individual data points corresponding to each distinct experimental community were systematically left out of sample, and models were then fit to all remaining data points using 10-fold cross-validation. The observed versus predicted values of left-out points estimated via this approach for a second-order model are shown in the scatter plots of Fig. 2. The prediction quality (*R*^2^) for left-out points are shown in white bars in Fig. 2.

### Calculation of Fourier amplitude spectrum

Because Eqn. 1 with *x*_*i*_ *∈* {*−*1, 1} corresponds to the Fourier expansion of a fitness landscape [30], the Fourier amplitude equation [31] at order *p* can be written simply as

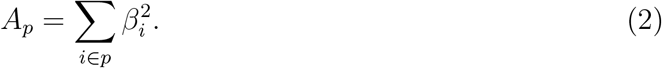

It can be shown that sum of amplitudes across all orders is equal to the total variance of the landscape. This permits the calculation of the fraction of total variance explained by terms at order *p* as *A*_*p*_*/* ∑_*p′*_ *A*_*p′*_.

### Calculation of *r/s* ruggedness metric

The roughness-slope ratio, *r/s*, is a measure of landscape ruggedness that quantifies how well a landscape is fit by a purely additive model. This quantity was computed by first fitting a linear model of the form:

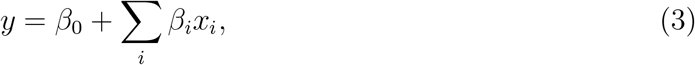

where *y* is the measured community function, and the coefficients *β*_0_ and *β*_*i*_ are obtained ordinary least-squares regression.

The roughness, *r*, is defined as the root-mean-squared-error of the resulting fit, or

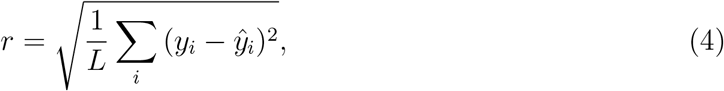

where *L* is the number of datapoints in the landscape and *ŷ* is the value of *y* fitted by linear regression. The slope, *s*, is defined as the mean of the absolute value of coefficients *β*_*i*_,

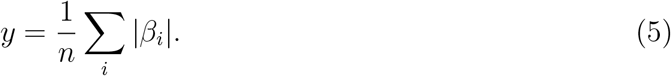

Larger values of *r/s* indicate a greater deviation from linearity, therefore a more rugged landscape. In contrast, an *r/s* value of 0 would correspond to a perfectly additive landscape. The values of *r/s* depend on landscape size, and sensible comparisons of this quantity between datasets requires an appropriate normalization. Because randomized landscapes represent a natural high-ruggedness comparison, *r/s* values computed on randomized landscapes were used as scaling factors, i.e,

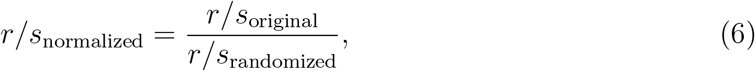

was computed. Normalized *r/s* values close to 1 indicate that the empirical landscape is nearly as rugged as the landscape randomization, while values close to zero indicate a low degree of ruggedness. 100 randomizations were performed for each dataset, and values of *r/s*_normalized_ were computed for each; the distributions of these values are shown in Fig. 3C.

### Simulations of the generalized Lotka-Volterra model

Community-function landscapes were synthetically generated using the generalized Lotka-Volterra (gLV) model, with total abundance serving as the community function of interest. A four-parameter “reference model” formulated by Barbier *et al*. [36] was used to explore important dimensions of the gLV parameter space. The model can be written as follows:

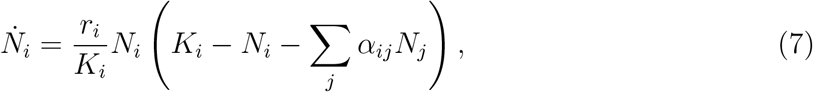

where, for species *i* in a pool of *N* total species, *N*_*i*_ is the abundance, *r*_*i*_ is the intrinsic growth rate, and *K*_*i*_ is the carrying capacity. The parameters *α*_*ij*_ are interaction coefficients between species *i* and *j*.

Nonzero equilibria of Eqn. 7 are determined by the values of *K*_*i*_ and *α*_*ij*_. For each synthetic landscape, these parameters were randomly generated. The carrying capacities *K*_*i*_ were independently drawn from a normal distribution with mean 1 and variance *ζ*^2^, while the interaction coefficients were drawn from a normal distribution with mean *µ/N* and variance *σ*^2^*/N*. A broad, ecologically-relevant region of the interaction parameter space was explored in Fig. 4A (*µ ∈* [*−*2, 4] and *σ ∈* [0, 1.5]). The standard deviation of carrying capacities was fixed at *ζ*= 0.3. In principle, interaction asymmetry can be encoded via a parameter *γ* = corr(*α*_*ij*_, *α*_*ji*_), but interactions here were set to be symmetric by taking *γ* = 1. All *r*_*i*_ were fixed to 1. These parameter values and ranges are identical to those used in Fig. S2 of Ref. [36], allowing direct comparison with Fig. 4A.

Landscapes were generated over a grid of points in *µ*-*σ* space. Each landscape was generated through the following steps:

1. A set of carrying capacity parameters *K*_*i*_ and interaction parameters *α*_*ij*_ were drawn for a pool of *N* = 10 species as described above.
2. Eqn. 7 was simulated to equilibrium for all species combinations.
3. Total endpoint abundances Σ _*i*_ *N*_*i*_(*∞*) were computed for each simulation.
4. Exact, full-order landscapes (Eqn. 1) were computed using total endpoint abundance as the community function.

Initial species abundances were drawn from an exponential distribution with mean 0.1. Numerical integration was performed using ode15s (MATLAB). At each point *µ*-*σ* space, 10 trials of landscapes were generated. Note that for larger values of *σ*, species abundances in Eqn. 7 are more likely to diverge. Parameter combinations for which divergences were encountered in more than half of trials are indicated by grey values in Fig. 4A.

### Simulations of the consumer-resource model with random cross-feeding networks

Community-function landscapes were synthetically generated using a consumer-resource model (CRM), taking the equilibrium concentration of a terminal waste product as the function of interest. The CRM is given as follows:

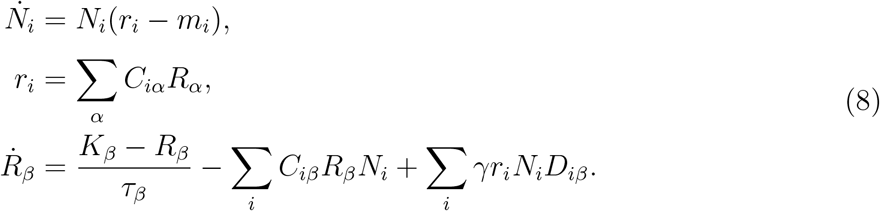

Here, *N*_*i*_, *r*_*i*_, and *m*_*i*_ are the abundance, total resource uptake, and maintenance costs, respectively, of species *i* in a pool of *N* total species. *R*_*α*_ is the concentration of resource *α*. The matrices *C*_*iα*_ and *D*_*iβ*_ describe which resources a species consumes and secretes, respectively, with secretions assumed proportional to the metabolic uptake *r*_*i*_. An efficiency factor *γ <* 1 ensures that energy cannot be gained, but only lost. Here *γ* was set to 0.5. The resource carrying capacities *K*_*β*_, decay rates *τ*_*β*_ and species maintenance costs *m*_*i*_ are all set to 1 for simplicity. The total number of resources was fixed to *L* + 1, with resource *L* + 1 representing a terminal waste product that no species can consume.

Random cross-feeding networks were generated in order to explore how chains of trophic dependencies impact the ruggedness of community-function landscapes. To do this, it was assumed that each species *i* can consume and secrete exactly one resource, denoted in_*i*_ and out_*i*_, respectively. These were selected through the following steps, performed independently for each species:

1. The identity of the consumed resource in_*i*_ was chosen randomly between 1 and *L* with equal probability.
2. The secreted resource out_*i*_ can range from 1 to *L*+1, where resource *L*+1 is the terminal waste product; however it must be distinct from in_*i*_, leaving *L* possible choices. With probability *p*, out_*i*_ = in_*i*_ + 1; otherwise out_*i*_ was drawn from any of the remaining values at random.

The parameter *p* thus allows the exploration of a range of network topologies, from long linear pathways with a high degree of trophic dependency (at *p* ≈ 1) to random graphs (at *p* ≈ 1*/L*).

After selecting in_*i*_ and out_*i*_ for all species, it was verified that resource *L*+1 is “reachable” through the network from resource 1, i.e., whether there exists at least one path from resource 1 to resource *L* + 1. If this was not the case, the functional landscape (the concentration of resource *L* + 1 for a given set of species) would be identically zero; such networks were discarded as invalid and the steps above were repeated until a valid circuit was obtained.

For Fig. 4B, random cross-feeding networks were generated by carrying out the steps above across a range of values of *p*, fixing *N* = 10 total species and *L* = 8 total resources. 20 valid random trials over 15 values of *p ∈* [1*/L*, 1) were generated in order to create 300 total random cross-feeding networks. The community-function landscapes for these networks were then computed by simulating Eqn. 8 to equilibrium for all species combinations (ode15s, MATLAB), taking the equilibrium concentration of resource *L*+1 as the function of interest, and fitting Eqn. 1 exactly. Initial species abundances were set to 0.1, and initial resource concentrations were set to *R*_*β*_(0) = *K*_*β*_.

## I. SUPPLEMENTARY MATERIALS

**TABLE S1.** Information about datasets used in this study.

| Function | # species | # communities | Ref. |
| --- | --- | --- | --- |
| Butyrate production | 25 | 577 | Clark *et al.* [22] |
| Xylose oxidation | 6 | 63 | Langenheder *et al.* [27] |
| Starch hydrolysis | 6 | 53 | Sanchez-Gorostiaga *et al.* [21] |
| Siderophore production | 8 | 225 | Diaz-Colunga *et al.* [28] |
| Total biomass | 8 | 160 | Diaz-Colunga *et al.* [28] |
| Biomass of a single strain | 14 | 4239 | Kehe *et al.* [29] |

